# Eavesdropping and crosstalk between secreted quorum sensing peptide signals that regulate bacteriocin production in *Streptococcus pneumoniae*

**DOI:** 10.1101/087247

**Authors:** Eric L. Miller, Morten Kjos, Monica I. Abrudan, Ian S. Roberts, Jan-Willem Veening, Daniel E. Rozen

**Author notes:** These authors contributed equally to this work (Eric L. Miller, Morten Kjos).

## Abstract

Quorum sensing (QS), where bacteria secrete and respond to chemical signals to coordinate population-wide behaviors, has revealed that bacteria are highly social. Here, we investigate how diversity in QS signals and receptors can modify social interactions controlled by the QS system regulating bacteriocin secretion in *Streptococcus pneumoniae*, encoded by the *blp* operon (<u>b</u>acteriocin-<u>l</u>ike <u>p</u>eptide). Analysis of 4 096 pneumococcal genomes detected nine *blp* QS signals (BlpC) and five QS receptor groups (BlpH). Imperfect concordance between signals and receptors suggested widespread social interactions between cells, specifically eavesdropping (where cells respond to signals that they do not produce) and crosstalk (where cells produce signals that non-clones detect). This was confirmed *in vitro* by measuring the response of reporter strains containing each of six different *blp* QS receptors to cognate and non-cognate peptides. Assays between pneumococcal colonies grown adjacent to one another provided further evidence that crosstalk and eavesdropping occur at endogenous levels of signal secretion. Finally, simulations of QS strains producing bacteriocins revealed that eavesdropping can be evolutionarily beneficial even when the affinity for non-cognate signals is very weak. Our results highlight that social interactions can mediate intraspecific competition among bacteria and reveal that competitive interactions can be modified by polymorphic QS systems.

## Introduction

Quorum sensing (QS) is a mechanism of intercellular communication that allows bacterial populations to coordinately regulate gene expression in response to changes in population density. QS is controlled by the secretion and detection of diffusible signaling molecules that, at threshold concentrations, lead to increased signal secretion and the induction of coupled downstream pathways (Miller & Bassler 2001; Waters & Bassler 2005). By this process, QS ensures that metabolically costly products are only produced when this would benefit the bacterial population, i.e. when they are at high concentrations (Waters & Bassler 2005). QS systems are coordinated by the fact that cells simultaneously send and detect a specific signal (Bassler et al. 1997; Redfield 2002; Waters & Bassler 2005), a characteristic that increases the likelihood that QS functions as a private message between clonemates that share evolutionary interests (Crespi 2001; West et al. 2006; Schluter et al. 2016). However, although QS works as an effective means of gene regulation in the laboratory in single strain cultures, QS in nature may be less reliable because it is susceptible to signal eavesdropping (i.e. where a promiscuous QS receptor can detect a QS signal not produced by that genotype) and signal crosstalk (i.e. where a non-specific QS signal can activate QS receptors in genotypes that produce other QS signals) (Redfield 2002; Atkinson & Williams 2009). This variation in QS signals and QS signal detection is widespread in nature (Bouillaut et al. 2008; Swem et al. 2008; Ansaldi & Dubnau 2004; Ji et al. 1997) and distinct from well-studied cheater/cooperator dynamics (e.g. Jiricny et al. 2010; Strassmann & Queller 2011). For example, signal-blind bacteria that produce, but are incapable of responding to, QS signals can engage in signal crosstalk to manipulate the behavior of other cells, e.g. by inducing them to produce expensive public goods (Diggle et al. 2007). Crosstalk and eavesdropping can occur even if all cells within a population are otherwise phenotypically wild-type if (i) QS signals and receptors are polymorphic and (ii) signals can bind and activate more than one receptor variant. Here we examine these issues using the polymorphic QS system regulating bacteriocin production in the Gram-positive opportunistic pathogen *Streptococcus pneumoniae*, where QS is integral for mediating intraspecific competition.

To initiate infection, *S. pneumoniae* must successfully colonize the nasopharynx and then persist during subsequent colonization attempts from other strains. Commensal carriage of *S. pneumoniae* is widespread, affecting up to 88% of children worldwide (Regev-Yochay et al. 2004; Wyllie et al. 2014), and between 5-52% of individuals are co-colonized with multiple strains (Wyllie et al. 2014; Sauver et al. 2000; García-Rodríguez & Fresnadillo Martínez 2002; Brugger et al. 2010). Interactions between different strains during colonization are common and dynamic, and the rate of clonal turnover — where one strain displaces another — occurs on a timescale of days to months (Meats et al. 2003; Turner et al. 2012). Among the factors thought to mediate intraspecific competition among pneumococcal strains are small anti-microbial peptides with narrow target ranges called bacteriocins (Dawid et al. 2007), which are regulated by QS. The most diverse bacteriocins in *S. pneumoniae* are encoded by the *blp* (*b*acteriocin-*l*ike *p*eptides) locus (Lux et al. 2007; Dawid et al. 2007). We recently showed that the number of possible combinations of bacteriocins and immunity genes at this locus can extend into the trillions, although only several hundred combinations are actually observed (Miller et al. 2016). As with other Gram-positive peptide signals, the QS signal peptide (BlpC) regulating the *blp* operon is constitutively produced at low levels, but is auto-induced at high levels once a threshold concentration has been reached (Lux et al. 2007). Secreted BlpC binds to the extracellular domain of the membrane-bound histidine kinase BlpH, and upon binding the kinase phosphorylates the response regulator BlpR (Fig 1b; De Saizieu et al. 2000; Reichmann & Hakenbeck 2000) which initiates production of the *blp* bacteriocin and immunity genes and increases production of the BlpC signal (De Saizieu et al. 2000). *blpC* expression is also enhanced by the induction of pneumococcal competence, which is regulated by the paralogous *com* QS signaling system (Kjos et al. 2016). Both ABC transporters BlpAB (Håvarstein et al. 1995) and ComAB (Kjos et al. 2016; Wei-Yun et al. 2016) cleave the N-terminal, double-glycine leader sequence of BlpC before export of the mature peptide signal by the same transporters (Fig. 1b). Using QS to regulate secretion presumably ensures that Blp bacteriocins are only produced when there is a sufficiently high cell number to allow these anti-competitor toxins to reach effective concentrations.

**Figure 1.**
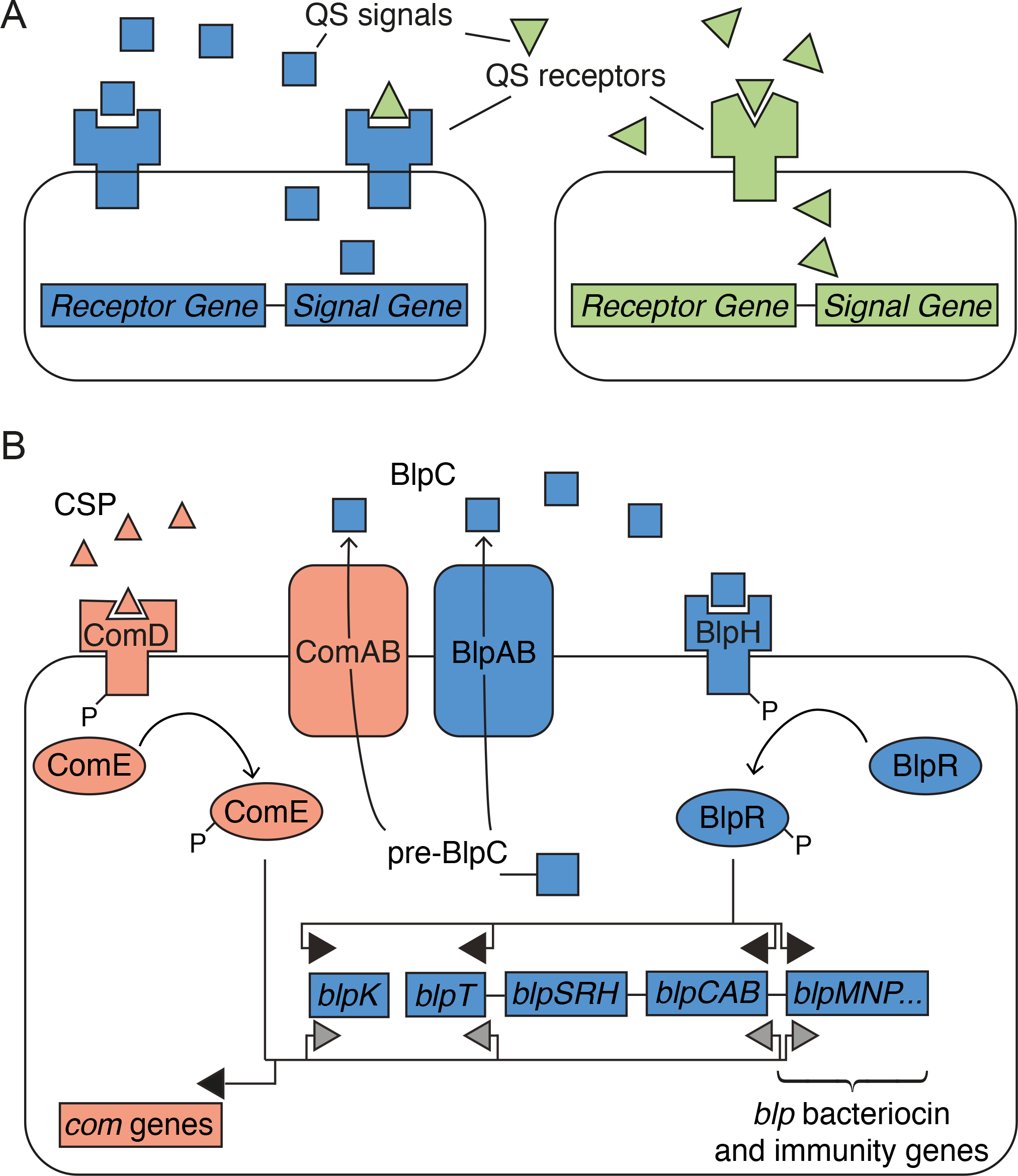
QS eavesdropping, crosstalk, and regulation. A) Eavesdropping occurs when a QS receptor of a cell is activated by a QS signal that the cell does not produce, such as activation of the blue QS receptor by both the cognate blue square signal and non-cognate green triangle signal. Crosstalk occurs when a QS signal activates more than one receptor, such as the green triangle signal activating both the cognate green QS receptor and the non-cognate blue QS receptor. B) *blp* QS regulation. External BlpC signal binds to histidine kinase receptor BlpH. This activates response regulator BlpR through phosphorylation, which increases transcription of *blpABC, blpT,* the *blp* bacteriocins (including *blpK*), and immunity genes. Pre-BlpC is processed and transported out of the cell by ABC transporters ComAB and BlpAB. Similarly, QS signal CSP binds to histidine kinase receptor ComD, thereby phosphorylating response regulator ComE, which increases transcription of the *blp* operon (although to a lower amount than BlpR) as well as *com*-specific genes.

Both the BlpC signal and its receptor, BlpH, are highly polymorphic (Miller et al. 2016). What are the effects of this variation, and how does this diversity influence the competitive interactions between strains that are mediated by *blp* bacteriocins? One possibility is that each unique BlpC signal corresponds to a distinct BlpH receptor to which it specifically and exclusively binds. By this explanation, strains detect and respond only to their own signal to determine the threshold at which they induce the *blp* operon. Such exclusivity is found in the *S. pneumoniae* competence signaling system where the two dominant peptide signals, CSP1 and CSP2, only induce cells expressing the cognate receptor (Iannelli et al. 2005). Similarly, there is near perfect concordance between the signal and receptor carried by any single genome, suggesting that tight coupling of these loci is crucial for the activation of competence (Miller et al. 2017). An alternative possibility, considered in a recent experimental study (Pinchas et al. 2015), is that BlpC peptides cross-react via crosstalk or eavesdropping with different BlpH receptors, thereby leading to a scenario where competing strains interact socially to induce the production of either immunity or bacteriocins at densities that would be insufficient for activation by auto-induction. Bacterial strains may benefit from this cross-reactivity if they are forewarned of the threats from others, allowing them to induce their own bacteriocins or immunity. Alternatively, eavesdropping may be costly if strains with promiscuous receptors are induced to secrete bacteriocins at densities that are too low to provide sufficient benefits to offset the costs of their production. *S. pneumoniae* presents an ideal opportunity to study the evolution of QS systems beyond cheater/cooperator dynamics (Pollak et al. 2016; Eldar 2011; Son et al. 2011) in an easily manipulated, highly relevant study system in which much is already known about signal/receptor dynamics (Pinchas et al. 2015). Our results reveal the importance of QS signaling polymorphism on *blp* operon regulation and clarify its ecological effects on *S. pneumoniae* intraspecific interactions.

## Materials and Methods

### Phylogenetic and Sequence Analysis

We analyzed *S. pneumoniae* genomes from eight publicly available sets, six of which contain strains that were randomly sampled from cases of disease or asymptomatic carriage: 3 017 genomes from refugees in Maela, Thailand (Chewapreecha et al. 2014); 616 genomes from Massachusetts carriage strains (Croucher, Finkelstein, et al. 2013); 295 genomes from GenBank, which include 121 genomes from Atlanta, Georgia, The United States (Chancey et al. 2015); 142 genomes from Rotterdam, the Netherlands (Hermans data set) (Bogaert et al. 2001; Miller et al. 2016); and 26 PMEN (Pneumococcal Molecular Epidemiology Network) genomes (McGee et al. 2001; Miller et al. 2016). The PMEN-1 (Croucher et al. 2011) and Clonal Complex 3 (Croucher, Mitchell, et al. 2013) data sets, containing 240 and 82 genomes, respectively, were a result of targeted sampling for specific clonal complexes of *S. pneumoniae*; as such, these strains were excluded from analyses that assumed random sampling. Details of the phylogenetic and sequence analysis are provided in the Supplemental Materials

### Bacterial strains and growth conditions

*S. pneumoniae* strains were grown as liquid cultures in C+Y medium (Moreno-Gamez et al. 2016) at 37°C and transformed as described previously (Kjos et al. 2016). For selection, *S. pneumoniae* was plated on Columbia agar supplemented with 2% defibrinated sheep blood (Johnny Rottier, Kloosterzande, Netherlands) and 1 μg/ml tetracycline, 100 μg/ml spectinomycin or 0.25 μg/ml erythromycin, when appropriate. *E. coli* was grown in LB medium with shaking at 37°C or plated on LA containing 100 μg/ml ampicillin.

#### Strain construction

Strains and plasmids used in this study are listed in Table S3. Full descriptions of strain construction for the expression of *blpSRH* alleles, the deletion of *blpSRHC,* and gene reporter constructs are given in the Supplemental Materials and Methods.

#### Luciferase assays

Luciferase assays were performed as described (Slager et al. 2014; Kjos et al. 2016). Briefly, *S. pneumoniae* cultures grown to OD_600_ 0.4 were diluted 100-fold in C+Y medium (pH 6.8) with 340 μg/ml luciferin. Luc-activity was measured in 96-well plates at 37°C, and OD_600_ and luminescence (as relative luminescence units, RLU) were recorded every 10 minutes using Tecan Infinite 200 PRO. Synthetic peptides (BlpCs) were purchased from Genscript (Piscataway, NJ). Different concentrations of BlpCs were added to the culture wells after 100 min or in the beginning of the experiment, depending on the experiment. The data was plotted as RLU/OD over time to analyze induction of *blp* expression.

### LacZ assays on agar plates

LacZ assays for testing induction by neighbouring colonies on plates were performed on C+Y agar (pH 8.0) covered with 40 μl of 40 mg/ml solution X-gal (spread on top of the plates). All strains were pre-grown to OD_600_ 0.4, before 2 μl of the wild-type strains (BlpC producers) were spotted and allowed to dry. Then 2 μl of the different reporter strains were spotted next to the dried spot. The plates were incubated at 37°C over-night.

For induction with synthetic BlpC, C+Y agar plates (pH 7.2) were covered with 40 μl of 40 mg/ml solution X-gal and 5 μl 1 mg/ml BlpC (spread on top of the plates), and different reporter strains were spotted on top. The plates were incubated at 37°C over-night.

### Stochastic Model

We built an individual-based spatial, stochastic model in which cells interact on a grid. We modeled four genotypes, which differ in the signaling molecule and bacteriocins that they produce as well as in the number and identity of signals that they respond to (Table S2). Bacteriocins produced by genotypes 1 and 2 specifically could kill genotypes 3 and 4 and vice versa. Signals produced by genotype 1 could induce genotypes 1 and 2 and similarly, signals produced by genotype 3 could induce genotypes 3 and 4; we therefore classify genotypes 2 and 4 as “eavesdropping genotypes”. Genotypes 1 and 3 can only respond to their own signal, as “signal-faithful genotypes”. All four genotypes have equivalent growth rates, which are only variable depending on if a cell is induced or uninduced. Eavesdropping cells respond to signals that they do not produce with certain degrees of affinity. If we consider the affinity of a cell to its own signal as 100%, we ranged the affinity to the other signals in the case of eavesdropping genotypes as 0% - 90% for different simulations. Full model details are given in the Supplemental Materials and Methods.

## Results

### Molecular diversity of *blpH* and *blpC*

We examined 4 096 *S. pneumoniae* genomes taken from six data sets of strains (Maela, Massachusetts Asymptomatic, GenBank, Hermans, Georgia GenBank, and PMEN: 4 096 genomes in total) alongside two additional data sets that are intentionally biased to specific clonal sub-groups (Complex 3 and PMEN-1: 322 genomes in total). We identified *blpC* in 99.0%, *blpH* in 99.0%, and both *blpC* and *blpH* in 98.2% of the combined 4 418 genomes using a DNA reciprocal BLAST algorithm (Miller et al. 2016). We note that the few genomes apparently lacking a *blp* gene may still contain these genes, as the data sets contain incomplete draft genomes. Consistent with earlier work (Miller et al. 2016), we found extensive allelic variation within *blpC,* which contains 37 alleles at the nucleotide level, 29 protein variants, and 20 different BlpC signal peptides, including signal peptides lacking a canonical double-glycine cleavage site. Nine of these peptide signal sequences were found in more than 0.5% of genomes (i.e., over 20 genomes; Table 1), and together these nine comprise ~98% of all signal variants. All signals under this 0.5% threshold were each confined to a single clade in the whole-genome phylogeny (Fig. S1). Each unique BlpC signal was designated with a letter from the NATO phonetic alphabet (Table 1). As expected for the genomes from intentionally biased samples, the PMEN-1 dataset almost exclusively carried the Golf signal (93.8%; Table S1), while the Clonal Complex 3 dataset almost exclusively carried the Kilo signal (97.6%; Table 1). The Bravo and Hotel signal peptides were exclusively found in strains collected as part of the Maela data set possibly indicating limited admixture between these strains and those from the other collections.

### *blpC/blpH* intragenomic pairing is highly biased but not exclusive

Phylogenetic analysis of *blpC* revealed four well-supported clades (Fig. 2) containing the following signals: 1) Alpha, Bravo, and Kilo; 2) Golf and Hotel; 3) Charlie; and 4) Delta, Echo, and Foxtrot. With the exception of the Delta signal, within-group signals are differentiated by a single amino acid or stop codon. The relationships between signaling groups within these major clades are uncertain, although there is evidence (0.75 < posterior probability < 0.95) that the Hotel, Bravo, and Delta signals are each monophyletic within their respective larger clades.

**Figure 2.**
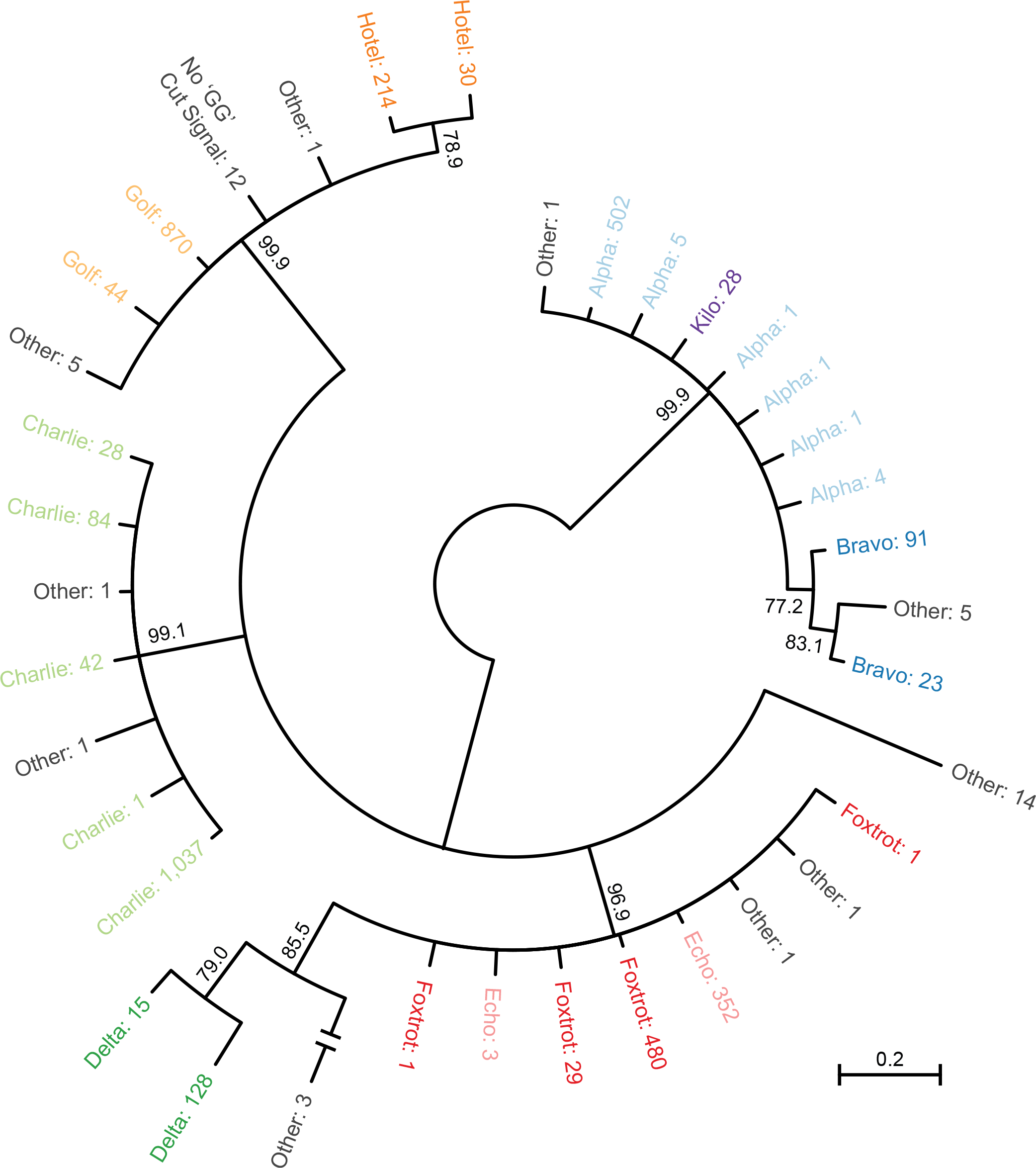
Bayesian unrooted phylogenetic tree of *blpC.* Taxa are colored by mature BlpC signal with the signal designation followed by the number of genomes containing the allele. Internal nodes show the posterior probabilities of clades; we collapsed clades with less than 0.75 posterior probability.

After accounting for recombination, phylogenetic analysis of the receptor domain of *blpH* (residues 1-229) identified five paraphyletic clades that are broadly concordant with the divisions observed for BlpC signals (Fig. 3), although there are many exceptions to this correspondence. Across the five clades, the classification of *blpH* alleles correlated with the BlpC signal in at least 75% of cases: (Alpha / Bravo / Kilo Clade: 86.6%; Echo / Foxtrot Clade: 90.0%; Delta Clade: 100%: Charlie Clade: 86.2%; Golf / Hotel Clade: 75.0%). Evidence of extensive recombination affecting the *blpH* kinase, intergenic region, and *blpC* signal (Fig. S2) suggests that recombination has caused some of these mismatches. Overall, from the 4 002 genomes with full-length *blpH* genes, 16.7% (667 genomes) show a lack of correspondence between signal and peptide, suggesting either that these strains are deficient in *blp* signaling or that these BlpH histidine kinase receptors can be cross-induced by non-cognate BlpC signals. Overall frequencies by signal and receptor class are summarized in Fig. 4a.

**Figure 3.**
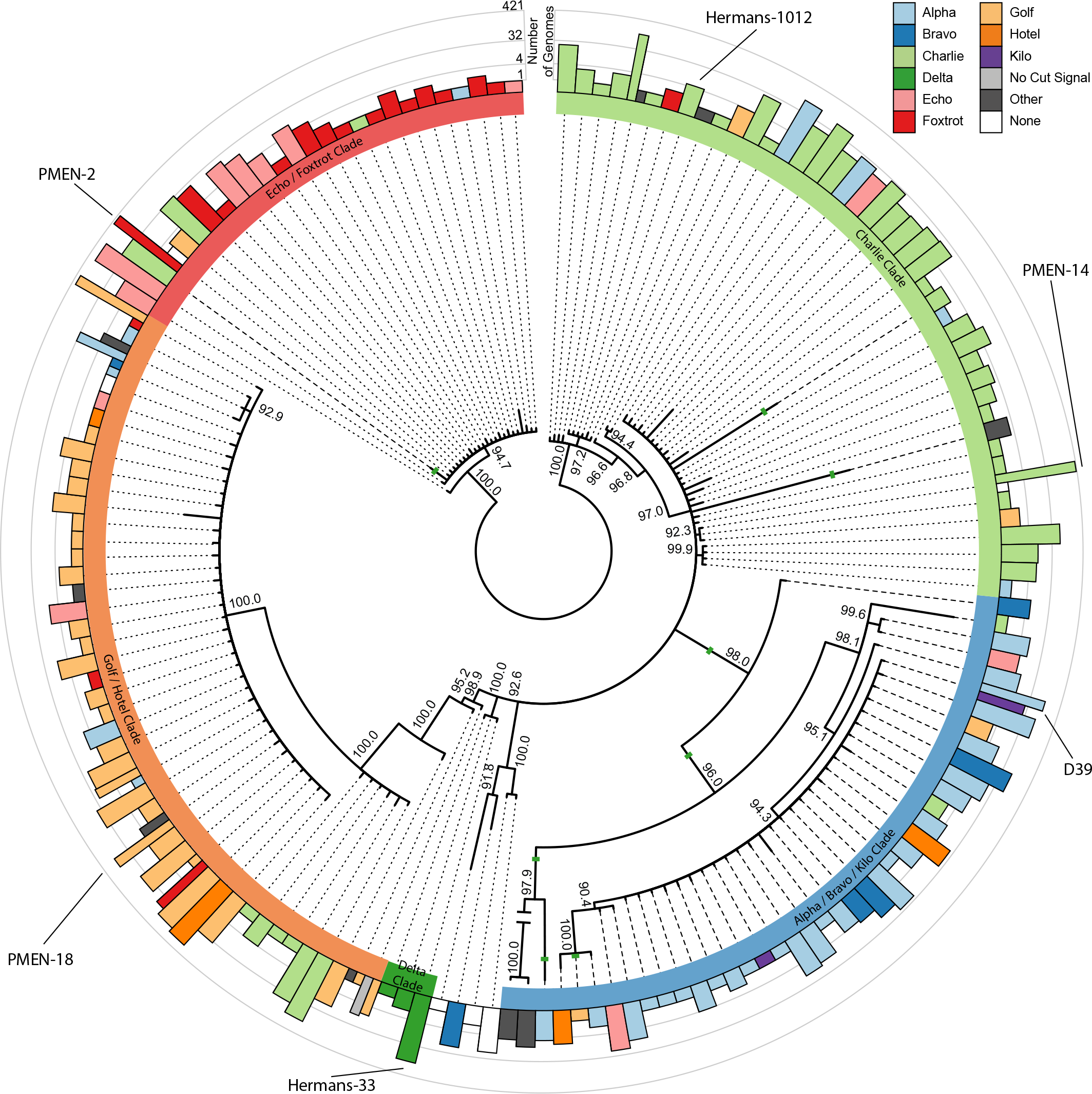
Bayesian unrooted phylogenetic tree of *blpH* alleles. The outer ring shows the number of 4 096 genomes with each *blpH* allele, color-coded by their co-occurring BlpC signal and on a log scale. The inner ring denotes the *blpH* clade type, and recombination events within *blpH* are shown as solid green lines. Mismatches between *blpH* clade and BlpC signal are indicated by dashed lines. Internal nodes show the posterior probabilities of clades; we collapsed clades with less than 90.0% posterior probability.

**Figure 4.**
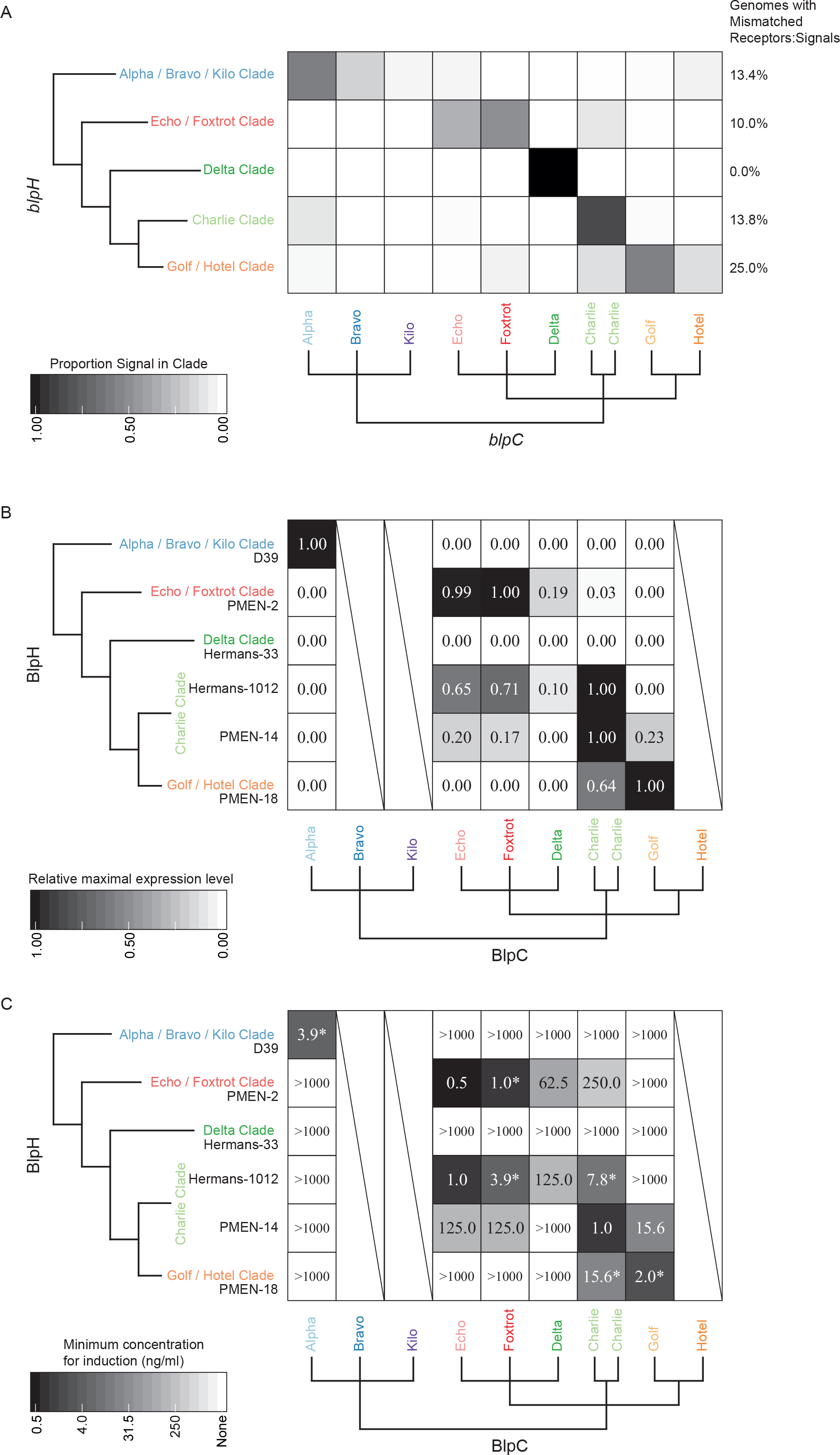
A) Proportion of each BlpC signal within genomes containing each *blpH* clade. The phylograms are simplified versions of Fig. 1 and Fig. 2. B) The relative maximal expression levels of *luc* following addition of 1 μg/ml of synthesized BlpC signal peptide. The maximum expression level for each reporter strain was set to 1. Raw data is found is Fig. S5 C) The minimum concentration of synthesized BlpC signal peptide required for *luc* induction in reporter strains with different BlpH. Asterisks indicate receptor/signal pair *blp* activation reported in (Pinchas et al. 2015). Example of raw data is provided in Fig. S6. The Bravo, Kilo, and Hotel signal peptides were not synthesized and are denoted with slashes.

### Crosstalk and eavesdropping between BlpC signals and BlpH receptors

To examine the incidence of crosstalk and eavesdropping between signals and receptors experimentally, we measured the responsiveness of each of the major BlpH clades to synthetic peptides from each BlpC class. We transformed a *S. pneumoniae* D39 strain lacking the native *blp* regulatory genes (*blpSRHC*) with constructs expressing one of six different BlpH histidine kinases alleles: *blpSRH*^D39^ from the Alpha/Bravo/Kilo clade, *blpSRH*^PMEN-2^ from the Echo/Foxtrot clade, *blpSRH*^Hermans-33^ from the Delta clade, *blpSRH*^Hermans-1012^ and *blpSRH*^PMEN-14^ from the Charlie clade, and *blpSRH*^PMEN-18^ from the Golf/Hotel clade. These strains also contained a reporter cassette, in which the *blp*-promoter from either P_*blpK*_ or P_*blpT*_ controlled expression of firefly luciferase (*luc*), GFP ( *(sf)gfp*), and β-galactosidase (*lacZ*) (Kjos et al. 2016). Deletion of the *blpC* signal gene and the native *blpSRH*genes from the D39 ancestor ensured that the reporter strains would only be induced in response to exogenously added signal via the introduced *blpSRH* systems. By exposing cells to a concentration gradient of exogenous peptide, we could estimate the peptide concentration that induced the maximum response as well the minimum concentration required to elicit a response. While the maximum response indicates the overall influence of a given peptide on each receptor, the minimal concentration required to induce a response provides an indication of the sensitivity of each receptor to every potential peptide partner.

Figures 4a-b shows that five of six P_*blpK*_ reporter strains were maximally induced by the BlpC signal carried by a significant majority of their wild type counterparts. However, we also see extensive evidence for crosstalk and eavesdropping between mismatched peptide:receptor pairs, demonstrating that some BlpH receptors are highly promiscuous while equally, several BlpC peptides can induce the *blp* operon in strains carrying non-complementary BlpH receptors. For example, *blpSRH*^PMEN-2^ (Echo / Foxtrot BlpH clade) could be induced by 4 out of 6 synthetic peptides, and the strain with *blpSRH*^Hermans-1012^ (Charlie BlpH clade) was strongly induced by the Echo and Foxtrot signals at 65% and 71% expression of its cognate signal. While there is clear evidence for cross-induction, these responses tended to be less sensitive to non-cognate peptides, with a minimum concentration required for induction of between 2–500–fold greater than with the cognate signal (Fig. 4c). By contrast, the strain with *blpSRH*^Hermans-1012^ (Charlie BlpH clade) was more sensitive to the non-cognate Echo and Foxtrot signals (1 ng/ml and 3.9 ng/ml) than to its complementary Charlie signal (7.8 ng/ml; Fig. 4c). The reporter strain carrying *blpSRH*^Hermans-33^ did not respond to any of the BlpC peptides, not even its cognate Delta BlpC (Fig. 4b-c). Interestingly, *blpSRH*^Hermans-33^, as well as all other strains with *blpH* alleles in the Delta clade, contains a frameshift in the *blpR* gene, encoding the response regulator, thus preventing expression of the full-length *blpR.* This probably renders the QS systems non-functional and therefore not responsive to added peptide. All results with P_*blpK*_ were mirrored with a different set of reporter strains that used the *blpT* promoter for the reporter cassette (Fig. S3).

We conclude from these results that crosstalk among quorum-dependent peptide BlpC signals is common and concentration dependent, with strains able to eavesdrop onto multiple signals using cross-responsive receptors. Furthermore, these results are concordant with the patterns of co-association observed in our bioinformatics survey of pneumococcal strains. When only considering genomes carrying *blpC* and *blpH* alleles potentially capable of *blp* activation (as determined in Fig. 4b and 4c), 88.0 % of the strains are predicted to autoinduce *blp* expression under appropriate conditions, i.e., their genomes contain functionally active *blpC/ blpH* pairs. Notably, however, this also indicates that a substantial proportion of strains (12.0%, 364 of 3 046 genomes) may not be able to autoinduce *blp* expression since they carry *blpC/blpH* pairs that were inactive in our experimental assay; this is in addition to strains carrying Delta *blpC/blpH,* which was also unable to autoinduce *blp* expression in our assay.

### Cross-induction between colonies

Pneumococci in the nasopharynx live in spatially structured colonies or biofilms. In order to determine if cross-induction between signaling cells could occur under these conditions where QS efficiency may be limited by signal diffusion (Redfield 2002; Kümmerli & Brown 2010; Yang et al. 2010), we examined interactions between neighboring colonies endogenously secreting either cognate or non-cognate signals (Fig. 5). In control assays, we first demonstrated that colonies were induced by exogenous addition of peptide to the plate surface; these results were concordant with those in Figure 4b in 14 of 15 combinations (Fig. S4). Next, we measured expression of reporter strains when grown adjacent to wild-type colonies that secreted BlpC peptides at endogenous levels (Fig. 5A). We observed a response in the reporter strains as estimated by increased LacZ activity in 3 out of 6 strains, with 2 examples of induction by noncognate BlpC signals. Interestingly, when the reporter strain expressing the BlpH from Hermans-1012 was grown adjacent to its wild type counterpart, there was no induction; instead this strain was induced by PMEN-14, which also produced the Charlie signal. The same strain was also induced by PMEN-2, which produced the Foxtrot signal (which induces Hermans-1012 at a lower concentration than with its cognate signal; Fig. 4C), and strain PMEN-18 (Golf/Hotel BlpH clade) was induced by PMEN-14, which produced the Charlie signal (Fig. 5). This may suggest that in addition to differences in the binding affinities of BlpC and BlpH, strains may also vary in the concentration of the diffusible signals that they secrete, at least under these experimental conditions. Consistent with our *in vitro* assays with synthesized peptides, these results show that *blp* operon expression can be activated by crosstalk between neighboring competing colonies secreting peptides at wild-type concentrations.

**Figure 5.**
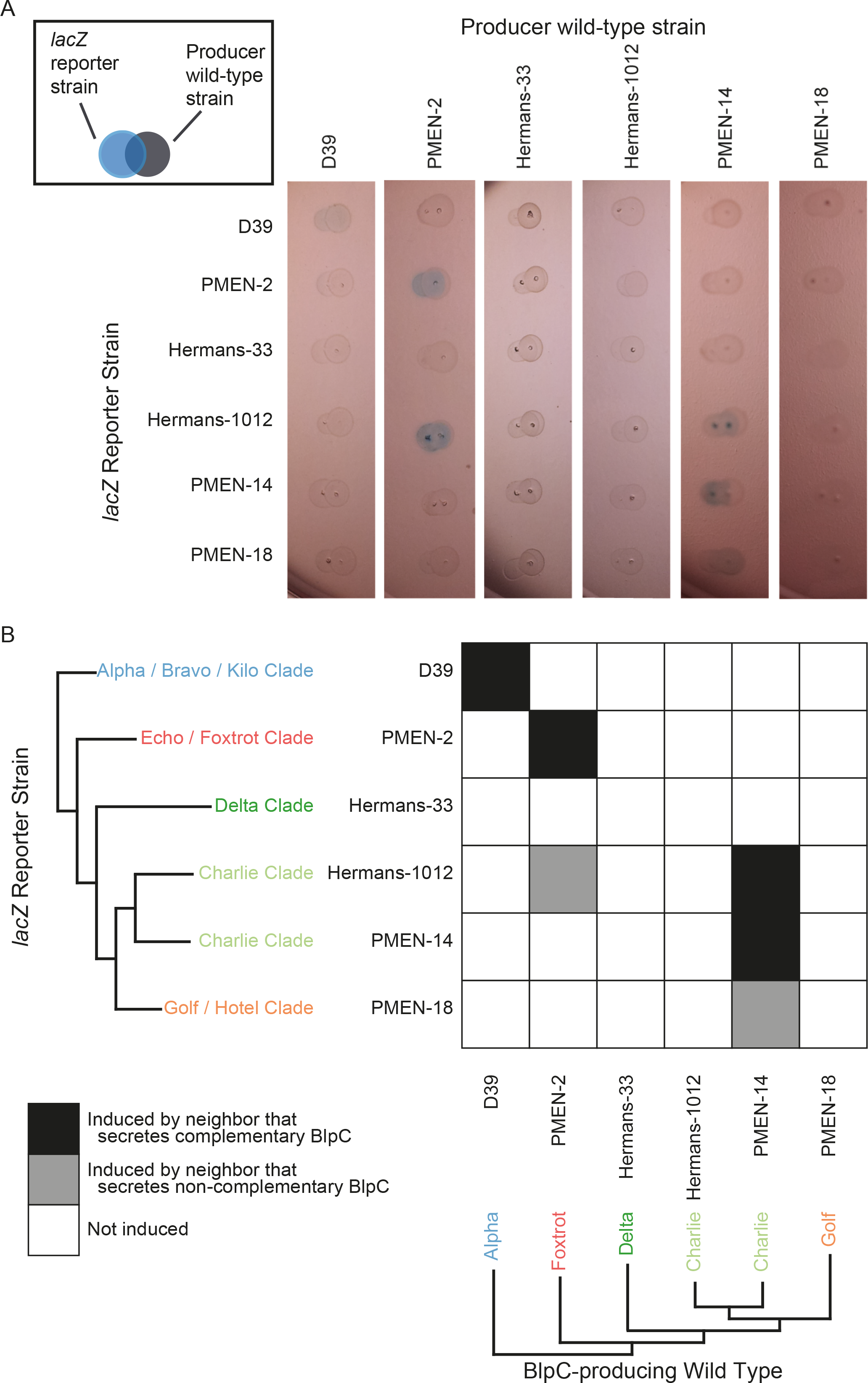
LacZ induction by neighboring colonies on agar plates. A) The wild-type strains were spotted next to the reporter strains (see box), and induction of *blp* expression by the wild-type produced BlpC is shown as faint blue colonies. The experiment was repeated three times with the same result, and a representative photo of the plates is shown. B) Summary of the results from B. Squares in white indicate no induction of the reporter strain for colony pairs, while black and blue indicate induction by complementary and on-complementary BlpCs, respectively.

### Evolutionary consequences of eavesdropping genotypes

Because the *blp* operon is auto-induced via a quorum-dependent process, cross-induction can potentially influence other strains by lowering the population density required for auto-induction. To examine the possible effects of cross-induction on bacteriocins, we developed a spatially explicit stochastic model to investigate conditions where genotypes with eavesdropping receptors may be favored over strains only able to respond to a single peptide signal. We further varied the signal affinity to eavesdropping receptors to determine how this altered the selective benefits of cross-induction. Simulations are initiated with cells of four genotypes randomly spaced upon a plane. The four genotypes each release their own QS signal at equal concentrations (Table S2). Cells bind these secreted signals in a concentration dependent manner, at which point they are induced to produce bacteriocins that kill susceptible neighbor cells at the cost of reduced growth for the producer (Ruparell et al. 2016). While two faithful-signaling genotypes are only able to respond to their own signals, the two other eavesdropping genotypes can respond to multiple signals. Our results shown in Fig. 6 lead to two conclusions. First, we observe strong benefits to eavesdropping cells that depends on the degree of cross-sensitivity, or affinity, to non-cognate signals. Specifically, we found that higher affinity to non-cognate signal provides stronger ecological benefits. This results from earlier potential activation (Fig. S3) and secretion of bacteriocins in these cells, an effect that increases with greater affinity to non-cognate signals. Second, we find that the benefits to eavesdropping are strongly negative frequency-dependent, i.e. eavesdropping cells only gain benefits (in the form of earlier bacteriocin induction) when surrounded by faithful-signaling cells. When eavesdropping cells are rare, they benefit through maximum exposure to the alternative peptide, while after they increase in frequency they must rely solely on auto-induction. Because the benefits of eavesdropping are frequency-dependent, these simple simulations thus suggest that promiscuous receptor mutants with increased affinity to non-cognate signals will be able to rapidly invade populations of cells that can only respond to a single signal. Interestingly, the simulations also clarify that the affinity to non-cognate signals can be extremely low — even at 10% of the affinity to cognate signals — to provide benefits (Fig. 6).

**Figure 6.**
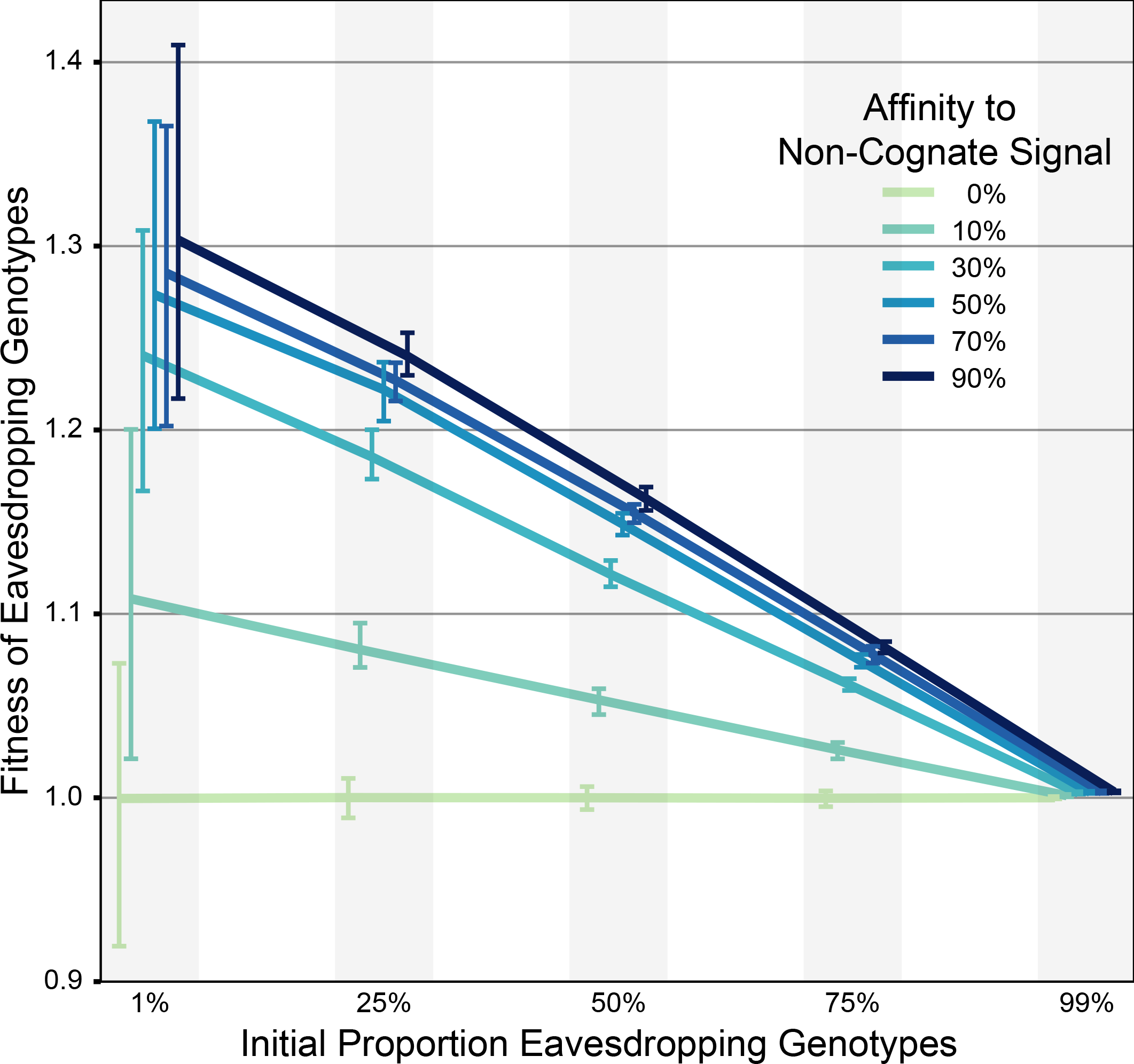
Average fitness of eavesdropping genotypes that produce bacteriocins in response to multiple signals in a spatially explicit, stochastic model. Simulations were started with five proportions of eavesdropping genotypes mixed with signal-faithful genotypes, as indicated on the x-axis. Absolute fitness values on the y-axis above 1.0 indicate that the genotype can increase in frequency in the population. Affinity to other genotypes’ signals are a percentage of affinity to a genotype’s own signal for eavesdropping genotypes. Error bars link the 25% and 75% quantiles for the final eavesdropping genotypes’ fitness across 100 simulations.

## Discussion

Pneumococcal bacteriocins are believed to play an important role in mediating intraspecific competitive interactions (Dawid et al. 2007). Here, we show that the QS system regulating *blp* bacteriocins is highly polymorphic, that QS signals are frequently cross-reactive (crosstalk), and that promiscuous receptors can detect and respond to noncognate signals (eavesdropping). Assays between adjacent colonies revealed that both behaviors occur at endogenous concentrations of secreted peptides, and simulations showed ecological benefits to strains that express promiscuous receptors. Together, these results suggest that social interactions influenced by QS signaling may strongly influence pneumococcal competition.

Previous surveys (De Saizieu et al. 2000; Reichmann & Hakenbeck 2000) of BlpC and BlpH identified four BlpC signals: the Alpha, Charlie, Foxtrot, and Golf signals in our nomenclature, which together represent ~75% of the strains in our sample (Table 1). By expanding our survey to thousands of strains, we identified several additional signal peptide families (Fig. 2): the Echo, Hotel, Delta, Bravo, and Kilo signals. The concordance between the phylogenies of *blpC* and *blpH* and the extensive co-occurrence in individual genomes suggest that these genes are co-evolving (Fig. 2, Fig. 3).

While the correlation between *blpH* clade and co-occurring BlpC signal is high, in some clades the correlation drops to 75.0%, and BlpH / BlpC mismatches (Fig. 3) are common across the pneumococcal phylogeny. This can be compared to the exceptionally tight, > 99% correlation between the ComD QS receptor and CSP signal also in *S. pneumoniae* (Miller et al. 2017). There are at least two explanations for this difference. First, we do not know if different BlpH variants are functionally distinct. All *blpH* alleles could, in principle, be most responsive to their cooccurring BlpC. This seems unlikely, given the high frequency (up to 45 signal:receptor pairs) of *blpH* clade / BlpC mismatches (Fig. 3). Second, weaker selection for a highly auto-inducing *blp* QS could explain the difference between the *blp* and *com* QS systems. After a recombination event that results in a sub-optimal BlpH/ BlpC pair for autoinduction, the BlpC signal may still be able to activate the co-occurring BlpH variant through crosstalk, albeit at a higher concentration of BlpC (Fig. 4C). While auto-induction may be decreased, such a genotype would gain an eavesdropping receptor that can potentially detect signals of surrounding genotypes. For comparison, there is no eavesdropping between CSP pherotypes in the *com* QS system, and very rare signal/receptor mismatches (Iannelli et al. 2005; Miller et al. 2017).

Signal/receptor mismatches can result in two outcomes for cell:cell communication. First, cells may be unable to detect the signal that they produce, rendering them unable to auto-induce. The lack of QS activation in strains producing the Delta signal (Hermans-33; Fig. 4) seemingly fits this description; however, interestingly, this is not caused by signal / receptor mismatch because there is perfect concordance between the Delta signal and the Delta *blpH* clade, and no tested signal activated strains with Delta *blpH*. Instead, all 143 strains carrying the Delta signal have a frameshift in *blpR*, which suggests functional deterioration of the QS system in these strains, which has not yet led to deterioration of *blpH* and *blpC* These Delta BlpC strains are not simply ‘cheater’ cells, as they potentially continue to pay the cost of synthesizing BlpC if the *blpC* gene is actively transcribed. This suggests there may be weakened selection for functional *blp* QS.

The second outcome of signal/receptor mismatches for cell-to-cell communication is crosstalk and eavesdropping. We have ample evidence for crosstalk in the *blp* QS system, as all signal peptides except for the Alpha signal activated QS receptors in genotypes that produce other QS signals (Fig. 4b-c). Similarly, BlpH receptors (aside from the Alpha clade) were eavesdropping QS receptors able to detect more than one QS peptide signal (Fig. 4b, Fig. 4c). Each of the receptors we tested (except for the signal-blind BlpH Delta clade) was maximally induced with a single set of related signals and decreased to 3-71% with signals that the receptors were eavesdropping upon (Fig 4b). This suggests that there are no ‘generalist’ receptors that are able to listen to multiple signals with equal affinity. Crosstalk was observed in previous research on the *blp* system (Pinchas et al. 2015; see asterisks in Fig. 4c and Table 1 aternative signal names), and results from this study indicated that *blpH*alleles with more crosstalk were less sensitive to BlpC (Pinchas et al. 2015). However, the results reported here show that receptors from strains PMEN-2, Hermans-1012, and PMEN-14 were all highly sensitive to their complementary signal (≤1.0 ng/ml) despite showing extensive crosstalk (Fig. 4c), thereby suggesting that the trade-off between crosstalk and sensitivity of *blpH*alleles is not a general phenomenon.

What are the potential consequences of crosstalk and eavesdropping? Crosstalk may enable one strain to manipulate competing strains into inducing their QS system at lower densities, thereby causing them to secrete bacteriocins and induced immunity proteins earlier. At present, it is unclear how such crosstalk would be beneficial to cells producing cross-reactive signals, unless premature production of bacteriocins or immunity introduces energetic or other costs to cells responding at sub-quorum densities. Similar benefits are thought to exist for other bacterial public goods (West et al. 2012; Diggle et al. 2007). By contrast, it is easier to envision the potential benefits of eavesdropping, which can both lead to earlier activation of bacteriocins (although this may also have attendant costs) and earlier induction of cross-reactive immunity. Our simulations suggest that this could be beneficial even if the affinity of promiscuous receptors is only 10% of the affinity for their cognate signal (Fig. 6). This value falls within the range of responses we measured experimentally (Fig. 4c). This level of responsiveness is also sufficient to induce the *blp* operon among adjacent colonies secreting peptides at endogenous levels (Fig. 5).

How does this amount of crosstalk specifically affect bacteriocin-mediated competition between *S. pneumoniae* strains? This is challenging to answer conclusively. First, extensive variation in the kinase domain of BlpH, the response regulator BlpR, and the leader sequences of the *blp* bacteriocins (Miller et al. 2016) prevents a full understanding of how signal concentrations translate into increased bacteriocin export. A systematic approach to investigate each of these molecules and their variants in the laboratory will be required to address this question. Second, a bioinformatics approach to examine evidence of selection in coordination with the BlpH receptor or BlpC signal is not possible due to the inability to align the entire *blp* operon and because recombination breaks up potential associations that are otherwise selected for. Third, the effects of crosstalk and eavesdropping will also depend on the activation of the *com* QS system, which promotes the expression and export of *blpC* at a low level (Fig. 1), even when the ABC-transporter genes *blpAB* are disrupted by early termination mutations (Kjos et al. 2016; Wei-Yun et al. 2016). For example, we found that both wildtype strains D39 and PMEN-14 could activate *blp* expression in neighboring colonies (Fig. 5) despite having disrupted *blpA* (for PMEN-14) or disrupted *blpA* and *blpB* (for D39).

Signaling interactions *in vitro* can lead to complex ecological outcomes that may influence competitive interactions between strains. As yet, however, it is unclear how these interactions will play out in the complex within-host environment of the human nasopharynx (Valente et al. 2016). In addition, it remains unclear how these interactions directly influence bacteriocin-mediated killing and immunity. Clearly, the heterogeneous conditions *in vivo* differ markedly between liquid cultures or agar surfaces. Diffusion is more limited, while population densities may be strongly constrained overall and spatially. These factors, among others, may alter the level and dispersion of signal peptides as well as the sensitivity of individual strains to these signals. More generally, our results reinforce the importance of social interactions among bacteria for mediating competitive dynamics. Many ecologically relevant bacterial traits are regulated by QS, and many of these systems, especially in Gram-positive peptide signaling systems, are polymorphic. While some of these systems (e.g. pneumococcal competence regulated by the *com* QS system) have only few signal types and show no cross-reactivity, many others signal are polymorphic with substantial cross-reactivity (e.g. Agr in *S. aureus* (Ji et al. 1997) and ComX in *B. subtilis* (Stefanic et al. 2012)). It remains to be investigated which of these polymorphic QS signals have ecological effects and which factors (such as co-colonization or extensive intraspecific competition) result in the evolution of crosstalk and eavesdropping.

## Acknowledgements

We would like to thank Frank Lake for technical assistance. This work was supported by the Biotechnology and Biological Sciences Research Council (grant number BB/J006009/1) to DER and ISR and by the Wellcome Trust (105610/Z/14/Z) to the University of Manchester. MA is supported by the Biotechnology and Biological Sciences Research Council (grant number BB/M000281/1). Work in the Veening lab is supported by the EMBO Young Investigator Program, a VIDI fellowship (864.12.001) from the Netherlands Organisation for Scientific Research, Earth and Life Sciences (NWO-ALW) and ERC starting grant 337399-PneumoCell. MK is supported by a grant from The Research Council of Norway (250976/F20).

